# Beyond Imbalance: An Elasticity Framework for the Distance-averaged Force-Velocity Relationship in Vertical Jump

**DOI:** 10.64898/2026.09.01.748530

**Authors:** Zhaoqian Li, Mingyu Li, Xing Zhang, Zongwei Chen, Litong Yang, Qiaozhe Li

**Author notes:** **Corresponding author:** Qiaozhe Li.

## Abstract

This study aimed to (1) establish the distance-averaged F-V relationship framework and (2) develop elasticity metrics that quantify how F-V relationship variables govern jump height and inform training prescription. Theoretical derivation and experimental validation across 108 F-V relationship models derived from 1578 jumps (countermovement jump and squat jump at three knee angles; 20 well-trained subjects) yielded a standard error of 2.1% and a nearly perfect correlation (r = 0.96, p < 0.001) between measured and predicted jump height. Four elasticity metrics were formulated: force elasticity (*F*_*e*_), the elasticity of jump height to maximal force (*F*_0_); velocity elasticity (*v*_*e*_), the elasticity of jump height to maximal velocity (*v*_0_); the force-velocity elasticity norm 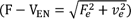, reflecting the overall sensitivity of jump height to changes in F-V relationship variables; and the force-velocity elasticity ratio (F − V_ER_ = *F*_*e*_ ÷ *v*_*e*_), indicating which variable dominates the jump height response. Simulations and experiments revealed that *F*_*e*_ bore an inverse relationship to *F*_0_, and *v*_*e*_ was inversely related to *v*_0_, reflecting diminishing marginal returns. At a fixed jump height, simulations showed F − V_EN_ and F − V_ER_ displayed a U-shaped relationship; a balanced profile (F − V_ER_=1) did not always correspond to the lowest F − V_EN_. The distance-averaged F-V elasticity framework offers a physically grounded and quantitative tool for linking F-V relationship variables directly to jump performance, providing a basis for informing individualized training decisions.

## Introduction

Vertical jump height is a fundamental indicator of athletic success and is frequently used to assess lower-limb neuromuscular function (Claudino et al., 2017; García-Ramos et al., 2021). Currently, the force-velocity (F-V) relationship is recognized as the most precise method for characterizing jump ability and designing individualized training direction (Li et al., 2026). This framework identifies two primary variables that constrain jump height: maximal power output (P_max_) and the F-V imbalance (F − V_IMB_) (Morin & Samozino, 2016). P_max_ is a strong predictor of jump height, while F − V_IMB_ determines whether an athlete’s unloaded jump is mainly limited by a force-based or velocity-based neuromuscular deficit (Morin et al., 2019). Athletes can therefore individualize their training based on F − V_IMB_ to accelerate improvement (Jiménez-Reyes et al., 2017). However, their modelling procedure’s assumption that distance-averaged velocity (*v̅*_*distance*_) equals time-averaged velocity has come under increasing scrutiny, since the former is usually higher than the latter (Bobbert et al., 2023; Bobbert et al., 2025; Linthorne, 2021). Given the inherent consistency observed in the force-time characteristics of vertical jumps, it is possible to leverage a physics-based determination of *v̅*_*distance*_ to address the validity concerns surrounding the F-V relationship (Linthorne, 2001). Such an approach would therefore offer a more robust framework for generalizing across diverse testing conditions.

However, even with an accurate distance-averaged framework, current F-V relationship training guidance does not yet rigorously quantify how the magnitude of F − V_IMB_ translates into jump performance, because under distance-averaged conditions, the product of mean force (*F̅*_*distance*_) and *v̅*_*distance*_ does not equate to mean mechanical power, precluding direct estimation of P_max_. To address this, given that jump height can be expressed as a quadratic function of the individual athlete’s *F*_0_ and *v*_0_, it may be possible to resolve the issue from an elasticity-based perspective. To formalize this concept, we define these sensitivity metrics as the elasticities of jump height with respect to *F*_0_ and *v*_0_, denoted as force elasticity (*F*_*e*_) and velocity elasticity (*v*_*e*_), respectively. These metrics quantify the percentage change in jump height resulting from a percentage change in *F*_0_ or *v*_0_, respectively. Practically, the ratio of *F*_*e*_ to *v*_*e*_, named F-V elasticity ratio (F − V_ER_) quantifies the relative sensitivity of jump height to *F*_0_ versus *v*_0_, providing a direct quantitative measure of the jump height response that complements the traditional F − V_IMB_, with implications for individualized training decisions. Additionally, the composite metric F − V elasticity norm (F − V_EN_), defined as the square root of the sum of squared *F*_*e*_ and *v*_*e*_, captures the combined jump height sensitivity to proportional changes in F-V relationship variables. Systematically investigating how these four elasticity metrics are modulated by underlying F-V relationship variables and characterizing their interdependencies may reveal whether they provide orthogonal or redundant information. These analyses also elucidate whether the metrics jointly inform training prescription beyond traditional method based on F − V_IMB_.

Accordingly, this study pursues two primary objectives: (1) to derive a theoretical solution for *v̅*_*distance*_, and to validate its accuracy for predicting jump height across countermovement jump (CMJ) and squat jump (SJ) at varying knee angles; (2) to elucidate how distance-averaged F-V relationship variables modulate the proposed elasticity metrics, and to characterize their internal interdependencies, through integrated theoretical derivation and experimental validation. We hypothesized that (1) the theoretically derived *v̅*_*distance*_ and distance-averaged F-V relationship would predict jump height with strong agreement to measured values pooled across CMJ and SJ at all three knee angles; and (2) *F*_0_ and *v*_0_ would modulate the four elasticity metrics through distinct patterns, with systematic interdependencies among the elasticity metrics.

### Theoretical Framework

#### Derivation of the Distance-Averaged Framework

Dynamic analysis of the vertical jump, combined with the chain rule, yields

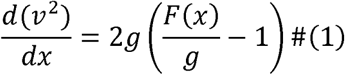

where *F*(*x*) is the ground reaction force normalized by body mass, and *g* denotes gravitational acceleration (9.81 m·s⁻²). Normalizing the ground reaction force and center-of-mass displacement as *f*(*x*) = *F*(*x*)/*g* and *u* = *x*/ℎ_*po*_, where ℎ_*po*_ is the vertical displacement of the center of mass from movement onset to take-off, and substituting into Equation (1), yields

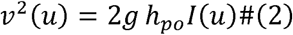

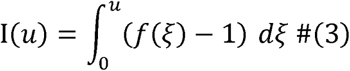

where I(*u*) is the cumulative net-work integral, representing the total propulsive impulse accumulated up to displacement *u*. The distance-averaged velocity follows as

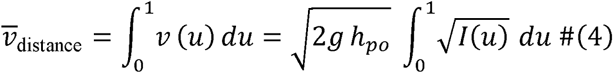

The take-off velocity follows as

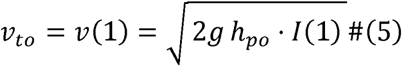

Forming the ratio of the distance-averaged velocity to the take-off velocity, the factor 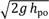 cancels, yielding

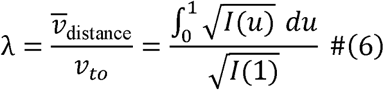

Crucially, λ depends only on the shape of *I*(*u*), independent of body mass, gravity, or ℎ_*po*_. Through analytical model and empirical calibration against force-time profiles and data from previous studies (Linthorne, 2001), λ converges to 0.77. Detailed derivation and the determination of λ are provided in Appendix 1. Since ℎ follows the projectile relation (Samozino et al., 2010):

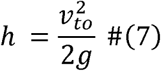

hence:

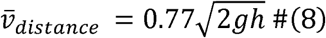

The mass-normalized *F̅*_*distance*_ has been extensively validated in prior studies, based on the work done during the unloaded vertical jump, and is given by equation (9) (Janicijevic et al., 2020; Samozino et al., 2008):

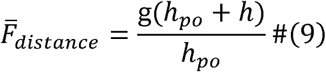

Here, the force-producing and velocity-producing capabilities of the lower limbs can be characterized by a linear force-velocity relationship (Pérez-Castilla et al., 2024; Rivière et al., 2023), where *F*_0_ (normalized by body mass) and *v*_0_ are the intercepts:

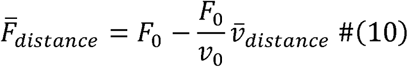

Incorporating equation (8) and equation (9) into equation (10), ℎ can be solved as:

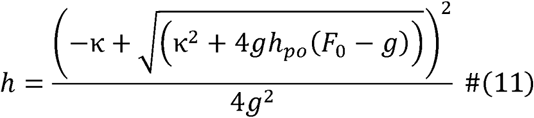

with 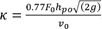.

#### Elasticity Metrics and Balance Condition

To intuitively characterize the effect of percentage changes in *F*_0_ and *v*_0_ on ℎ within the distance-averaged F-V relationship, the *F*_*e*_ and *v*_*e*_ were defined in equation (12) and (13):

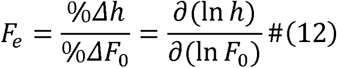

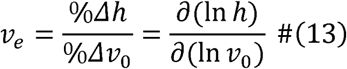

We define F − V_ER_ as the ratio of *F*_*e*_ to *v*_*e*_ in equation (14), and F − V_EN_ as the square root of the sum of the squares of the two elasticities in equation (15).

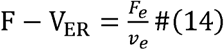

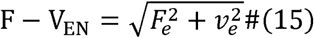

When F − V_ER_ = 1, it represents the traditional F-V balance condition; see Appendix 2 for the proof. Under this condition:

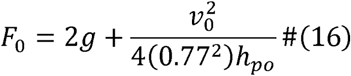

Taking the differential of both sides and simplifying yield:

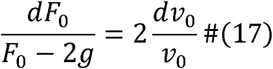

This relation states that, at the balance point, the fractional increase in (*F*_0_ − 2*g*) must be exactly twice the fractional increase in *v*_0_. Notably, this proportionality is independent of ℎ_*po*_. The detailed derivation is provided in Appendix 3.

## Method

### Numerical Simulation

According to equation (11) and previous studies, ℎ is determined by *F*_0_ (range from 20 to 60 N·kg⁻¹), *v*_0_ (range from 3 to 8m·s⁻¹), and ℎ_*po*_ (range from 0.2 to 0.6m). To systematically isolate the effects of each F-V relationship variable on the elasticity metrics, three sequential univariate scanning protocols were implemented: (1) modulating *F*_0_ across its range at seven fixed *v*_0_ levels (from 3.5 to 6.5 m·s⁻¹) with ℎ_*po*_ held at 0.40 m; (2) symmetrically varying *v*_0_ across its range at seven fixed *F*_0_ levels (from 25 to 55 N·kg⁻¹) under identical ℎ_*po*_; and (3) evaluating the sensitivity to ℎ_*po*_ using six representative (*F*_0_, *v*_0_) dyads spanning the experimental spectrum. *F*_*e*_ and *v*_*e*_ were computed from the scanning grid, and their complementary variation was examined. The composite metrics F − V_ER_ and F − V_EN_ were then constructed from *F*_*e*_ and *v*_*e*_, and the relationship between them was examined across the full grid under the identical ℎ and ℎ_*po*_. The condition *F*_*e*_ = *v*_*e*_ was subsequently identified as a special case within this relationship, corresponding to F − V_ER_ = 1.

### Participants

Twenty male well-trained volunteers (age: 22.0 ± 1.3 years; body mass: 77.0 ± 7.8 kg) participated in this study. All participants had at least one year of resistance training experience and were free from any lower-limb or musculoskeletal injuries within six months prior to the study. All participants were familiar with performing unloaded and loaded CMJ and SJ at various knee angles. The study was approved by the local ethics committee and conducted in accordance with the Declaration of Helsinki. Written informed consent was obtained from all participants prior to participation.

### Experimental Protocol

Participants performed CMJs and SJs under three knee-angle conditions: shallow (70°-80°), standard (≈ 90°), and deep (100°-110°). The six testing conditions (3 knee angles × 2 jump types) were randomized across separate days, with each condition tested on a distinct day. On each testing day, participants performed jumps with external loads of 0, 20, 40, 60, and 80 kg using a barbell placed on the shoulders. Three trials were performed at each load, applied in ascending order. One minute of rest was allowed between trials at the same load, and three minutes between different loads. Prior to data collection, participants completed a standardized warm-up consisting of jogging, dynamic stretching, and several vertical jumps without and with a light external load. For the SJ, participants descended to the predetermined knee angle, held the position for approximately two to three seconds, and were instructed to jump as high as possible without any countermovement. For the CMJ, participants performed a natural countermovement to their target depth before jumping. Not all participants completed all six conditions. A total of 1578 jumps were performed, and 108 successful testing sessions were completed across all conditions.

### Equipment and Data Analysis

All jumps were performed on a Kunwei resistive force plate force platform (KWYP-FP6035, Kunwei, Shanghai, China) with the vertical ground reaction force sampled at 1000 Hz (Mao et al., 2024). For SJ, the onset of the concentric phase was defined as the instant when the ground reaction force exceeded baseline by 20 N (Pérez-Castilla et al., 2019). Take-off was identified as the instant when the force fell below 20 N (Merrigan et al., 2024). Jump height was calculated from the take-off velocity, determined by integrating the net force-time curve from onset to take-off. The push-off distance was obtained by double integration of the net force signal. For CMJ, the same integration was performed from the lowest center-of-mass position as onset to take-off. *F̅*_*distance*_ and *v̅*_*distance*_ were computed from the center-of-mass force-displacement curve. For each F-V relationship model, the jump with the highest jump height at each load was selected for modelling. A linear regression was fitted to the F-V relationship across loads to determine *F*_0_ (force intercept), *v*_0_ (velocity intercept) and P_max_ = 0.25*F*_0_*v*_0_ (Li et al., 2024). Predicted *v̅*_*distance*_ and jump height were calculated from equations (8) and (11).

### Statistical Analysis

Descriptive statistics are presented as mean ± SD. Normality of the data was verified using the Shapiro–Wilk test (p > 0.05). All analyses were performed using SPSS (version 25.0, IBM, Armonk, NY, USA) with statistical significance set at p < 0.05. All jumps were pooled across conditions and loads. The agreement between predicted and measured *v̅*_*distance*_ and jump height was evaluated using systematic error (SE = mean of prediction − measurement, with 95% confidence interval), random error (RE = SD of the differences), and the percentage counterparts SE% and RE% relative to the mean measured value (Morel et al., 2026). Pearson’s correlation coefficient (r) was used to assess the linear association between predicted and measured values. Pearson correlation coefficients were also calculated (1) between the F-V relationship variables and the elasticity metrics, (2) between *F*_*e*_ and *v*_*e*_ and (3) between F − V_ER_ and F − V_EN_. The magnitude of the r coefficient as: trivial (< 0.10), small (0.10-0.29), moderate (0.30-0.49), large (0.50-0.69), very large (0.70-0.89), and nearly perfect (≥ 0.90) (Z. Li et al., 2025).

## Results

### Validation of the distance-averaged force-velocity model

The measured *v̅*_*distance*_ was 1.620 ± 0.341 m·s⁻¹, predicted *v̅*_*distance*_ was 1.654 ± 0.368 m·s⁻¹, yielding a small systematic error of SE = 0.034 [0.030; 0.038] m·s⁻¹ (SE% = 2.1%), RE = 0.089 m·s⁻¹ (RE% = 5.5%), and a nearly perfect Pearson correlation of *r* = 0.97 (*p* < 0.001). The measured height was 0.411 ± 0.056 m, predicted height was 0.403 ± 0.056 m, with a small systematic error of SE = 0.008 [0.005; 0.012] m (SE% = 2.1%), RE = 0.017 m (RE% = 4.1%), and a nearly perfect Pearson correlation of *r* = 0.96 (*p* < 0.001).

### Effects of force-velocity relationship variables on elastic metrics

Three univariate scanning protocols were computed over the empirically representative parameter space and validated against all experimental observations. Simulation curves predicted that *F*_*e*_ decreases and *v*_*e*_ increases monotonically with rising *F*_0_, yielding declining F − V_ER_ and F − V_EN_. The experimental data confirmed this pattern: athletes with higher *F*_0_ exhibited substantially lower *F*_*e*_ (*r* = −0.94, *p* < 0.001) and higher *v*_*e*_ (*r* = 0.44, *p* < 0.001). Correspondingly, both F − V_EN_ (*r* = −0.68, *p* < 0.001) and F − V_ER_ (*r* = −0.64, *p* < 0.001) decreased with increasing *F*_0_ (Figure 1).

**Figure 1.**
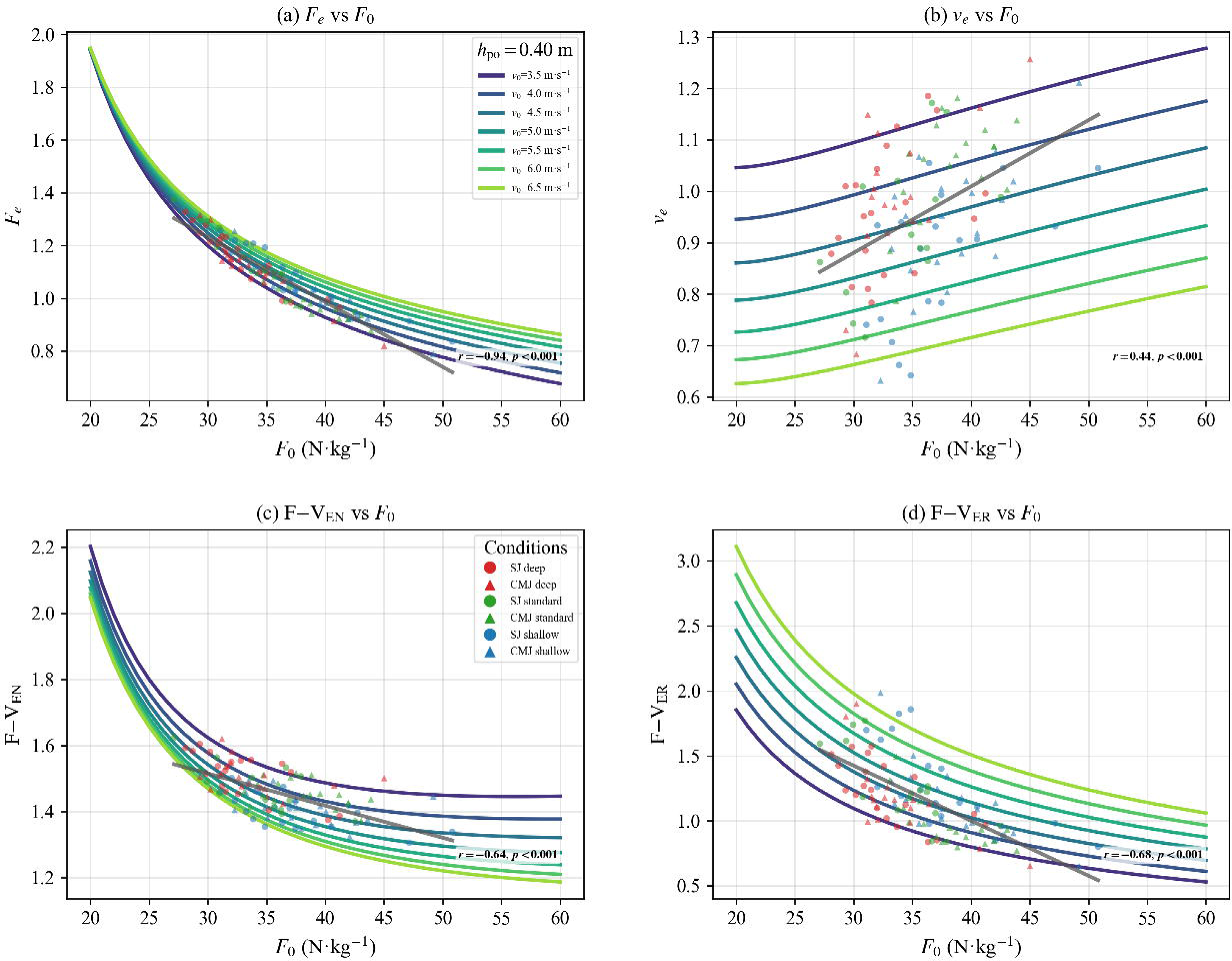
Influence of increasing maximal force ability (*F*_0_) on elasticity metrics derived from the distance-averaged F-V relationship: (a) force elasticity (*F*_*e*_), (b) velocity elasticity (*v*_*e*_), (c) force-velocity elasticity norm (F − V_EN_), and (d) force-velocity elasticity ratio (F − V_ER_). Data are presented for squat jumps (SJ, circles) and countermovement jumps (CMJ, triangles). Colors indicate knee angle conditions: red = deep, green = standard, blue = shallow. Bold indicates statistical significance.

Conversely, increasing *v*_0_ at fixed *F*_0_ shifted the simulated elasticity metrics toward higher *F*_*e*_ and lower *v*_*e*_, yielding a rise in F − V_ER_ and a decline in F − V_EN_. Experimentally, athletes with higher *v*_0_ exhibited higher *F*_*e*_ (*r* = 0.75, *p* < 0.001) and lower *v*_*e*_ (*r* = −0.92, *p* < 0.001), driving F − V_ER_ upward (*r* = 0.93, *p* < 0.001). A negative correlation was observed between F − V_EN_ and *v*_0_, but it was not statistically significant (*r* = −0.17, *p* = 0.085) (Figure 2).

**Figure 2.**
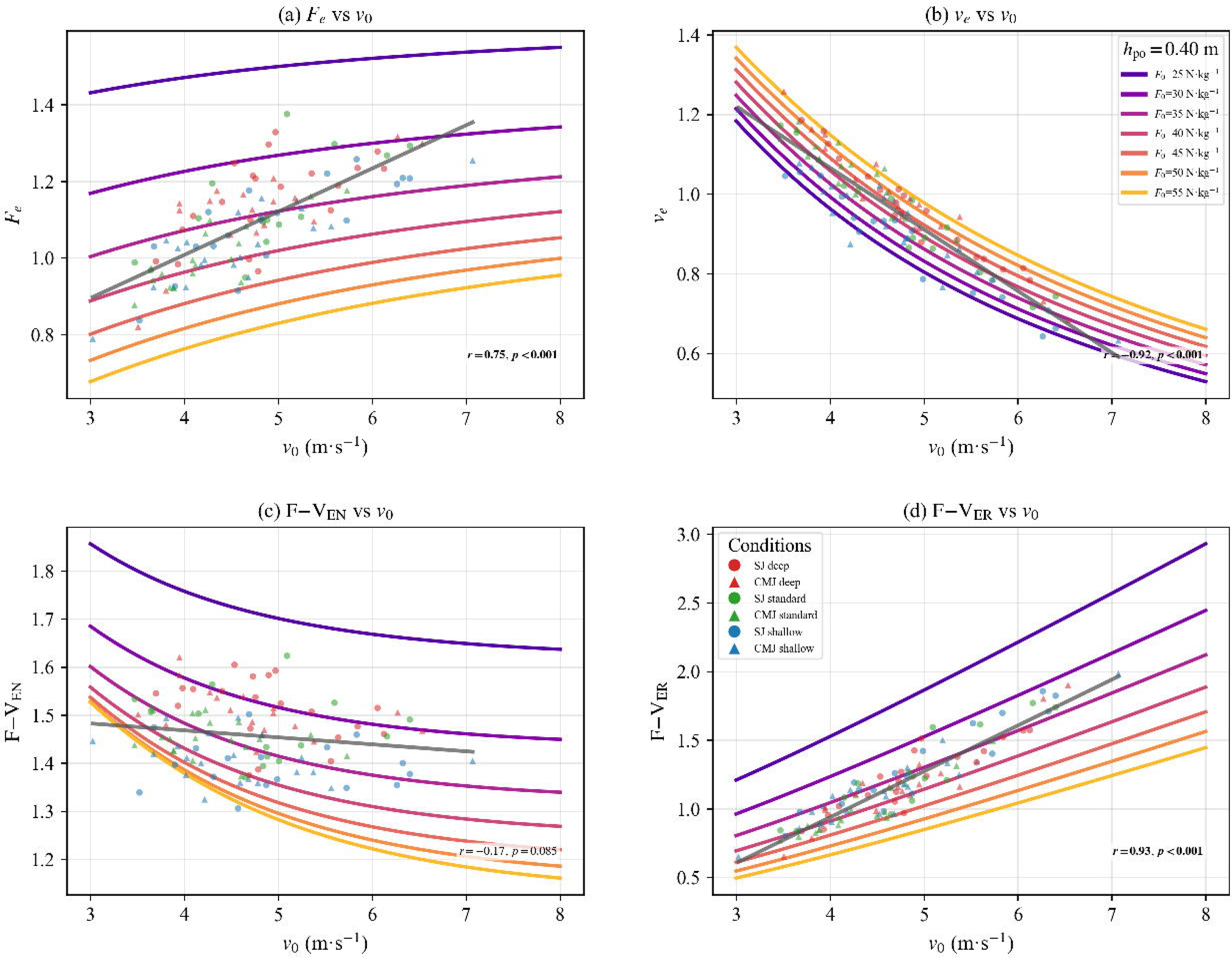
Influence of increasing maximal velocity ability (*v*_0_) on elasticity metrics derived from the distance-averaged F-V relationship: (a) force elasticity (*F*_*e*_), (b) velocity elasticity (*v*_*e*_), (c) force-velocity elasticity norm (F − V_EN_), and (d) force-velocity elasticity ratio (F − V_ER_). Data are presented for squat jumps (SJ, circles) and countermovement jumps (CMJ, triangles). Colors indicate knee angle conditions: red = deep, green = standard, blue = shallow. Bold indicates statistical significance.

**Figure 3.**
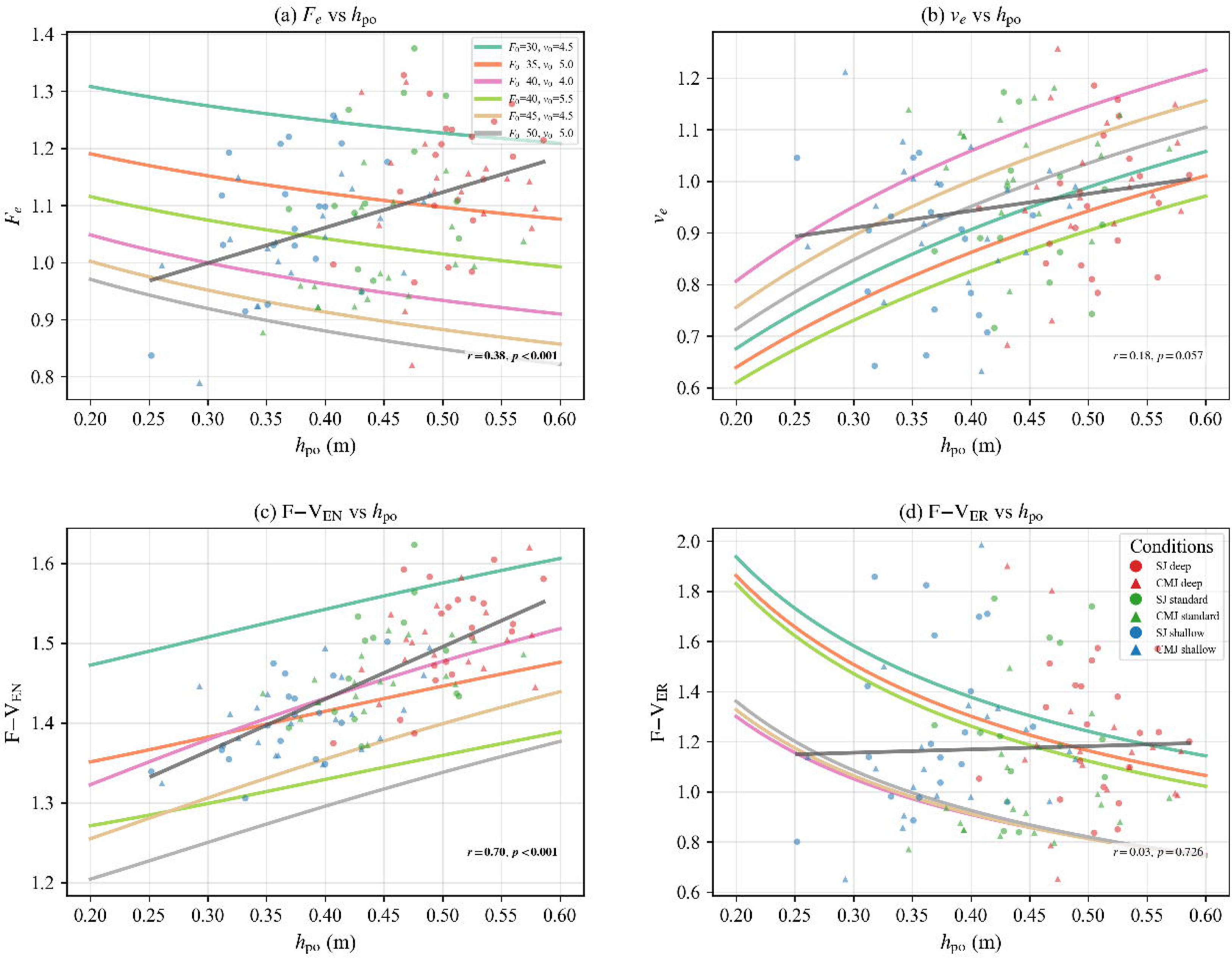
Influence of increasing push-off distance (ℎ_*po*_) on elasticity metrics derived from the distance-averaged F-V relationship: (a) force elasticity (*F*_*e*_), (b) velocity elasticity (*v*_*e*_), (c) force-velocity elasticity norm (F − V_EN_), and (d) force-velocity elasticity ratio (F − V_ER_). Data are presented for squat jumps (SJ, circles) and countermovement jumps (CMJ, triangles). Colors indicate knee angle conditions: red = deep, green = standard, blue = shallow. Bold indicates statistical significance.

The sensitivity scan over ℎ_*po*_ using six representative (*F*_0_, *v*_0_) pairs yielded the following predictions: a greater ℎ_*po*_ increases *v*_*e*_ but modestly decreases *F*_*e*_. Consequently, F − V_EN_ rises whereas F − V_ER_ falls. Experimental data only partially supported these predictions for F − V_EN_, which showed an increasing trend with ℎ_*po*_(*r* = 0.70, *p* < 0.001). Contrary to the predicted decrease, *F*_*e*_ exhibited a moderate positive correlation with ℎ_*po*_ (*r* = 0.38, *p* < 0.001). The corresponding trend for *v*_*e*_ was positive but not significant (*r* = 0.18, *p* = 0.057), and F − V_ER_ was uncorrelated with ℎ_*po*_ (*r* = 0.03, *p* = 0.726).

### Interplay between elasticity metrics

The simulated density, colored by ℎ_*po*_, revealed a continuous negative association between *F*_*e*_ and *v*_*e*_. Using the thresholds *F*_*e*_ = 1 and *v*_*e*_ = 1, four qualitatively distinct quadrants were identified: velocity-dominant (*F*_*e*_ < 1, *v*_*e*_ > 1), dual-high (*F*_*e*_ > 1, *v*_*e*_ > 1), dual-low (*F*_*e*_ < 1, *v*_*e*_ < 1), and force-dominant (*F*_*e*_ > 1, *v*_*e*_ < 1). This pattern was reproduced in the experimental data, with a significant negative correlation between *F*_*e*_ and *v*_*e*_ (*r* = −0.69, *p* < 0.001) (Figure 4).

**Figure 4.**
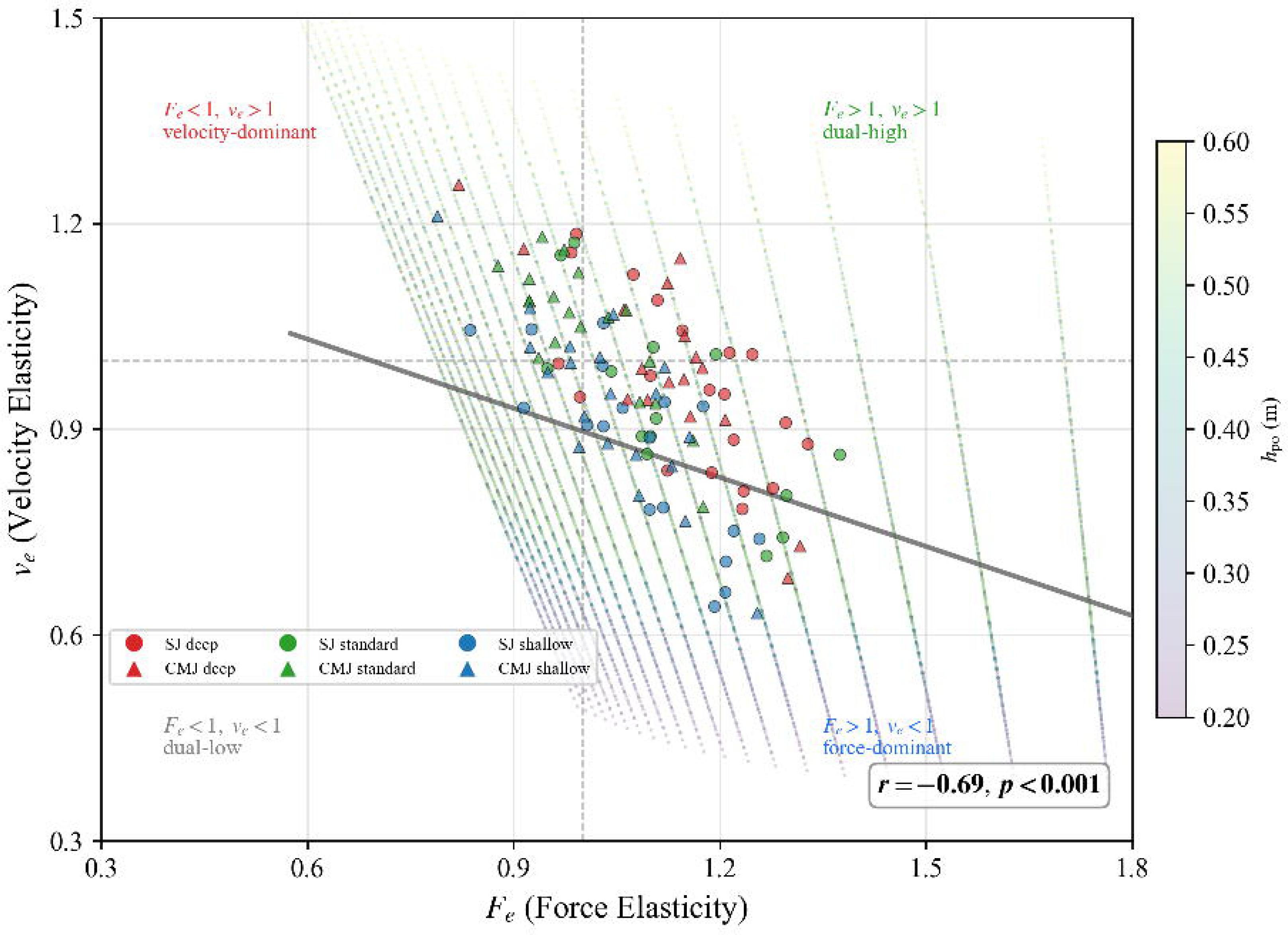
Negative association between force elasticity (*F*_*e*_) and velocity elasticity (*v*_*e*_), spanning four quadrants: velocity-dominant (*F*_*e*_ < 1, *v*_*e*_ > 1), dual high (*F*_*e*_ > 1, *v*_*e*_ > 1), dual low (*F*_*e*_ < 1, *v*_*e*_ < 1), and force-dominant (*F*_*e*_ > 1, *v*_*e*_ < 1). Bold indicates statistical significance. ℎ_*po*_: push-off distance.

Under a constant ℎ_*po*_, F − V_EN_ exhibited a U-shaped dependence on F − V_ER_ for each jump height. The balanced condition F − V_ER_ = 1 does not universally minimize F − V_EN_; instead, the F − V_EN_ at the balanced condition decreases with increasing target jump height. The ridge of minimum F − V_EN_ shifted from F − V_ER_ ≈ 1.3 at low jump height to F − V_ER_ ≈ 0.7 at high jump height (Figure 5). Experimental data showed no significant correlation between F − V_EN_ and F − V_ER_(*r* = −0.02, *p* = 0.868). Figure 6 depicts the required trade-off between the percentage changes in *F*_0_ and *v*_0_ needed to maintain the balanced condition (F − V_ER_ = 1) at different *F*_0_ levels: the higher the *F*_0_, the smaller the percentage change in *v*_0_ needed relative to that in *F*_0_. Specifically, the required ratio (*Δv*_0_/*v*_0_)/(*ΔF*_0_/*F*_0_) declines from ∼2.3 at *F*_0_ = 25 N·kg⁻¹ to ∼0.7 at *F*_0_ = 60 N·kg⁻¹.

**Figure 5.**
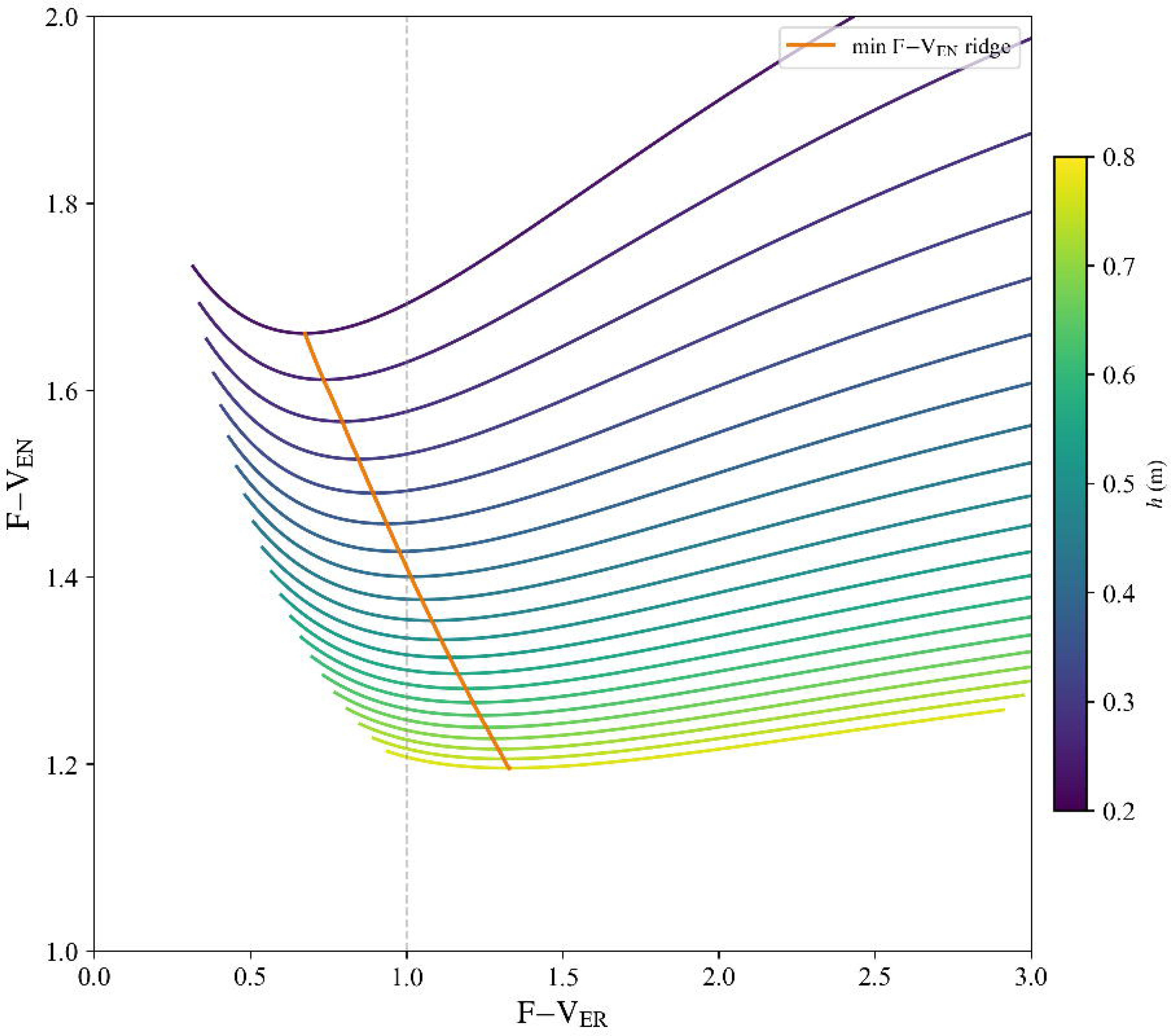
U-shaped relationship between F − V_EN_ and F − V_ER_ under constant push-off distance (ℎ_*po*_ = 0.4m). The yellow line shows the F − V_ER_ at which F − V_EN_ is minimized for each jump height.

**Figure 6.**
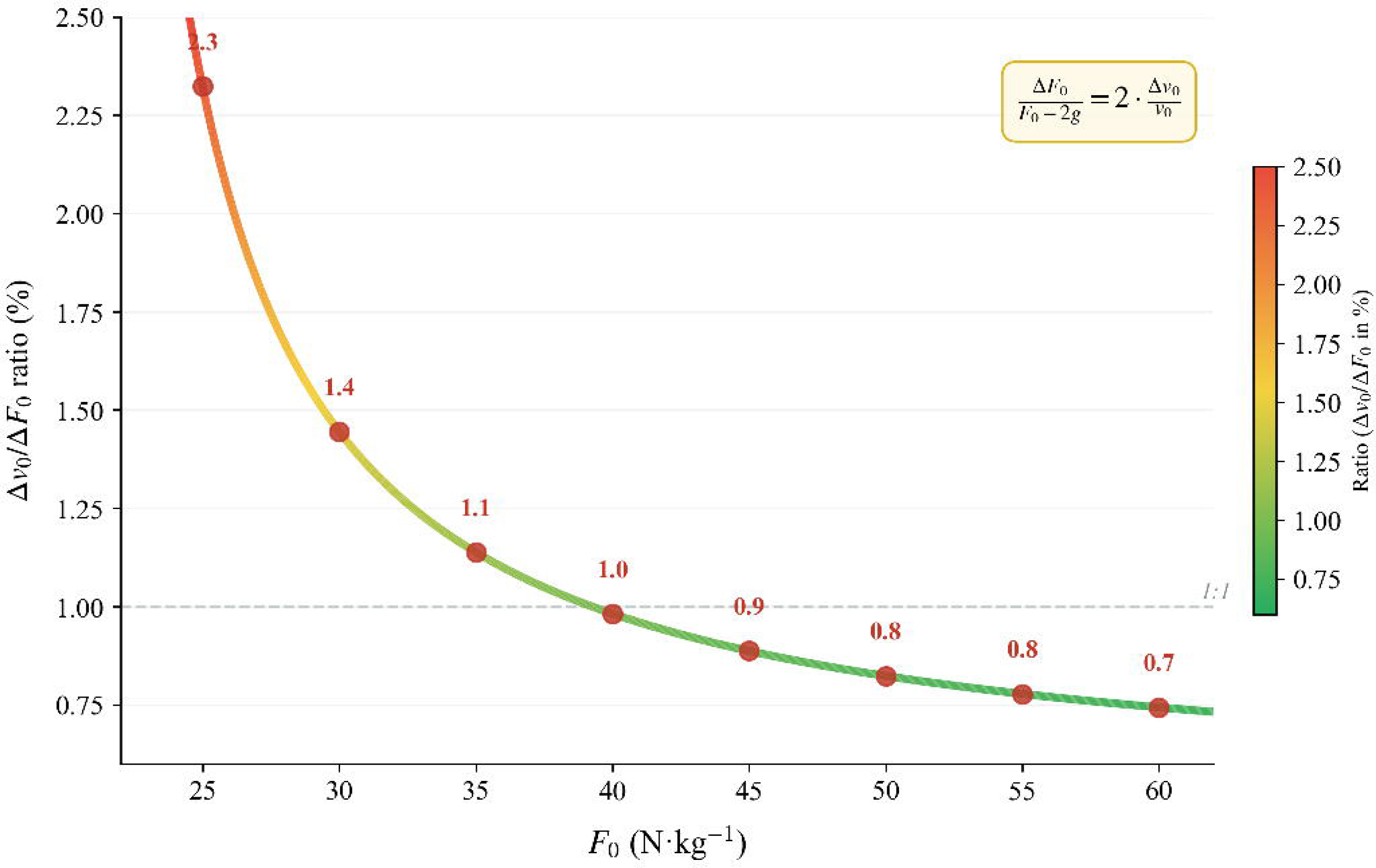
Percentage increase in maximal velocity ability (*v*_0_) required for a 1% increase in maximal force ability (*F*_0_) to maintain balance (F − V_ER_[force − velocity elasticity norm] = 1) across different baseline *F*_0_ levels.

## Discussion

The present study validates the distance-averaged F-V relationship model for predicting jump height and introduces an elasticity framework that quantifies the sensitivity of jump height to changes in F-V relationship variables. The principal findings were threefold. First, the distance-averaged F-V relationship model predicted *v̅*_*distance*_ and jump height with high accuracy, confirming its validity as a biomechanical framework. Second, *F*_0_ and *v*_0_ exerted opposing effects on the elasticity metrics: higher *F*_0_drove F − V_ER_ downward (via reduced *F*_*e*_ and elevated *v*_*e*_), whereas higher *v*_0_drove F − V_ER_ upward (via elevated *F*_*e*_ and reduced *v*_*e*_). Third, the balanced condition (F − V_ER_ = 1) did not coincide with the lowest F − V_EN_ at a given ℎ_*po*_ and jump height; experimental data showed no significant correlation between F − V_EN_ and F − V_ER_. Together, these results support the proposed elasticity framework as a valid approach for informing vertical jump training decisions. F − V_ER_ indicates which variable dominates the jump height response, while F − V_EN_ quantifies the combined jump height sensitivity to proportional changes in F-V relationship variables.

### Validation of the distance-averaged force-velocity model

Although a minor systematic bias was observed, with *v̅*_*distance*_ and jump height overestimated by 0.03 m/s and 0.01 m respectively, the measures demonstrated near-perfect agreement with prediction and minimal random variability. These findings confirm the validity of both the solution for *v̅*_*distance*_ and its predictive accuracy for jump height. Since *v̅*_*distance*_ depends on the shape of the force-displacement trajectory, direct calculation of *v̅*_*distance*_ requires additional assumptions about how force declines with center of mass displacement and the force at the lowest point (Linthorne, 2001). Through the double-integral analysis of Linthorne’s CMJ force-displacement profile, we revealed that λ is not a fixed constant but varies with both the rate of force decay and the force magnitude at the lowest position (Linthorne, 2021). The actual λ fluctuates within a modest range and clusters around 0.76 ± 0.05, differing only slightly from the value of 0.77 adopted here. In simplified methods, establishing F-V relationship variables does not require processing force-plate data. Instead, it relies solely on jump heights under different loads and measurements of the distance from the squatting position to triple extension, which serve as a proxy for ℎ_*po*_. The predictive equations for jump height and the derivation of mean velocity are based on the same underlying formulas in simplified method condition, the specific value of the coefficient λ does not affect prediction accuracy (Samozino et al., 2012). For instance, the traditional method of Samozino et al. employs a fixed coefficient of 0.5 (Samozino et al., 2014), the simplified methods yielded negligible systematic error and low random error.

### Effects of force-velocity relationship variables on elastic metrics

The present findings reveal a clear asymmetry in how the two intercepts of the F-V relationship influence the elasticity metrics in both simulation and experiment. The opposing effects of *F*_0_ and *v*_0_ on F − V_ER_ constitute the most consistent finding of this study. In both the simulated parameter scans and the experimental data, increasing *F*_0_ consistently reduced *F*_*e*_ and raised *v*_*e*_, driving F − V_ER_ downward; increasing *v*_0_ produced the mirror image, elevating *F*_*e*_ and suppressing *v*_*e*_, thereby driving F − V_ER_ upward. Increment along the F-V relationship in either direction tends to diminish the overall elasticity magnitude, while the direction of sensitivity is uniquely captured by F − V_ER_. Specifically, raising one intercept suppresses its corresponding elasticity, whereas it enhances the elasticity associated with the other intercept. Nevertheless, the overall F − V_EN_ consistently decreases with an increase in either intercept.

The effect of ℎ_*po*_ on the elasticity metrics was less consistent between the simulation and experiment. Although the simulation predicted that a greater ℎ_*po*_ would reduce *F*_*e*_ and elevate *v*_*e*_ (thereby lowering F − V_ER_ and raising F − V_EN_), only the increase in F − V_EN_ was experimentally confirmed. Notably, *F*_*e*_ showed a moderate positive correlation with ℎ_*po*_, opposite to the predicted decline. A likely explanation is the confounding between ℎ_*po*_ and *F*_0_ in the repeated-measures design: deeper squats increase ℎ_*po*_ but decrease *F*_0_ (Pommerell et al., 2025; Qin et al., 2025). Since a smaller *F*_0_ was associated with a larger *F*_*e*_, whereas the simulation predicted that a larger ℎ_*po*_ alone would reduce *F*_*e*_, the observed positive correlation suggests that the *F*_0_ driven increase in *F*_*e*_ outweighed the isolated negative effect of ℎ_*po*_. Therefore, the positive correlation between ℎ_*po*_ and *F*_*e*_ suggests that a smaller *F*_0_ had a stronger influence than the simulated ℎ_*po*_ effect.

#### Interplay between elasticity metrics

The complementary trade-off between *F*_*e*_ and *v*_*e*_ provides a mechanistic basis for the behavior of the elasticity metrics. In the simulation, F-V relationship variables projected onto the *F*_*e*_ and *v*_*e*_ plane revealed a dense negative association spanning four distinct quadrants: velocity-dominant (*F*_*e*_ < 1, *v*_*e*_ > 1), dual-high (*F*_*e*_ > 1, *v*_*e*_ > 1), dual-low (*F*_*e*_ < 1, *v*_*e*_ < 1), and force-dominant (*F*_*e*_ > 1, *v*_*e*_ < 1). This pattern was replicated in the experimental data (r = −0.69, p < 0.001). This negative coupling implies that athletes cannot simultaneously maximize both elasticities in the short-term, and training strategies must therefore prioritize one based on the athlete’s profile.

Under constant ℎ_*po*_, F − V_EN_ exhibited a U-shaped dependence on F − V_ER_ at each jump height, with the minimum value of F − V_EN_ ridge shifting from F − V_ER_ ≈ 1.3 at low jump height to ≈ 0.7 at high jump height. F − V_ER_ = 1 does not universally minimize the F − V_EN_. Instead, at higher jump demands the velocity-dominant profile (F − V_ER_ < 1) exhibits a progressively larger jump height response to changes in F-V relationship variables, whereas at lower jump heights the force-dominant profile (F − V_ER_ > 1) is more responsive. At the balance point where *F*_*e*_ = *v*_*e*_, the jump height responds with equal sensitivity to marginal changes in *F*_0_ and *v*_0_, and the percentage increase in *F*_0_ − 2*g* is always twice the percentage increase in *v*_0_. This relation is independent of ℎ_*po*_. Although the equality of *F*_*e*_ and *v*_*e*_ constitutes a mathematically well-defined condition, any athlete progressing from low to high *F*_0_ and *v*_0_ will inevitably pass through this balanced state. Rather than blindly pursuing balance as an end in itself, training should focus on a periodized exploration of whether to develop *F*_0_ or *v*_0_ first.

### Training-induced adaptations of F-V relationship variables

The elasticity metrics quantify how sensitively jump height responds to changes in *F*_0_ and *v*_0_, but this does not imply that training the variable with the larger elasticity will necessarily yield faster or greater improvement in F-V relationship variables and corresponding jump height. The magnitude of adaptation depends not only on the elasticity of the outcome but also on the trainability of each variable and the time course of specific interventions. The elasticity framework identifies the sensitivity of jump height; it does not prescribe which variable is more efficient to train. Both the athlete’s training history and the potential expected adaptive dose of each modality must be weighed when translating elastic diagnostics into training decisions.

Table 1 summarizes representative longitudinal studies that have compared different training strategies on F-V relationship variables, detailing how *F*_0_, *v*_0_, and P_max_ respond to each approach, including the rate of change per week (Barrera-Domínguez et al., 2023; Escobar Álvarez et al., 2020; Jiménez-Reyes et al., 2017; Jiménez-Reyes et al., 2019; Jonson et al., 2026; Simpson et al., 2021; Zabaloy et al., 2020). These data offer practical guidance for individualized training prescriptions. In most training conditions, P_max_ remains insensitive to targeted F-V relationship shifts or even decreased following optimized training protocols (Jiménez-Reyes et al., 2017; Jiménez-Reyes et al., 2019; Zabaloy et al., 2020). For instance, in the first phase of combined optimized training group reported by Barrera-Domínguez et al. (Barrera-Domínguez et al., 2023), P_max_ decreased by 6.69% despite clear shifts in *F*_0_ (+8.45%) and *v*_0_ (−17.82%). The only exception was the power training protocol in Jonson et al., where a substantial increase in P_max_ (+16.60%) was observed, driven primarily by a large gain in *v*_0_ (+15.42%) rather than *F*_0_ (+0.24%) (Jonson et al., 2026). Therefore, P_max_ should not be used as a standalone monitoring variable; instead, *F*_0_ and *v*_0_must be assessed independently to capture meaningful mechanical adaptations.

**Table 1.** Training adaptations and weekly progression rates of F-V relationship variables.

| Group | Study | Subjects | $\Delta P_{\max}$ | $\Delta P_{\max} \cdot wk^{-1}$ | $\Delta F_0$ | $\Delta F_0 \cdot wk^{-1}$ | $\Delta v_0$ | $\Delta v_0 \cdot wk^{-1}$ |
| --- | --- | --- | --- | --- | --- | --- | --- | --- |
| force-deficit training | Jiménez-Reyes et al. (2017) | athletes | -0.65% | -0.07% | 22.11% | 2.46% | -19.81% | -2.20% |
| force-deficit training | Escobar Álvarez et al. (2020) | ballet dancers | 0.09% | 0.01% | 24.18% | 2.69% | -20.00% | -2.22% |
| force-deficit training | Zabaloy et al. (2020) | rugby players | -5.44% | -0.78% | 10.90% | 1.56% | -14.80% | -2.11% |
| high force-deficit training | Jiménez-Reyes et al. (2019) | athletes | -3.16% | -0.20% | 43.12% | 2.71% | -33.33% | -2.10% |
| low force-deficit training | Jiménez-Reyes et al. (2017) | athletes | 3.53% | 0.38% | 16.41% | 1.78% | -10.97% | -1.19% |
| velocity-deficit training | Zabaloy et al. (2020) | rugby players | 3.96% | 0.57% | -13.74% | -1.96% | 17.65% | 1.96% |
| velocity-deficit training | Jiménez-Reyes et al. (2017) | athletes | 9.92% | 1.10% | -6.45% | -0.72% | 20.18% | 2.10% |
| high velocity-deficit training | Jiménez-Reyes et al. (2019) | athletes | 7.72% | 0.80% | -10.36% | -1.08% | 12.50% | 1.56% |
| low velocity-deficit training | Jiménez-Reyes et al. (2019) | athletes | 7.92% | 0.99% | -3.97% | -0.50% | 20.09% | 2.87% |
| Well-balanced training | Jiménez-Reyes et al. (2017) | athletes | 5.44% | 0.60% | 1.56% | 0.17% | 3.63% | 2.87% |
| Well-balanced training | Zabaloy et al. (2020) | rugby players | -3.08% | -0.44% | 3.48% | 0.50% | -5.32% | -0.76% |
| non-optimized | Jiménez-Reyes et al. (2017) | athletes | 2.13% | 0.24% | -2.64% | -0.29% | 4.17% | 0.46% |
| non-optimized | Simpson et al. (2021) | rugby players | 10.26% | 1.28% | -0.09% | -0.01% | 10.36% | 1.29% |
| non-optimized | Zabaloy et al. (2020) | rugby players | -0.35% | -0.05% | -0.56% | -0.08% | 1.49% | 0.21% |
| non-optimized | Barrera-Domínguez et al. (2020)* | basketball players | 5.00% | 1.25% | -1.56% | -0.39% | 9.79% | 2.45% |
| non-optimized | Barrera-Domínguez et al. (2020)** | basketball players | 5.49% | 1.37% | 0.63% | 0.16% | 3.34% | 0.84% |
| Optimized training mixed | Simpson et al. (2021) | rugby players | 3.82% | 0.48% | 0.81% | 0.10% | 2.99% | 0.37% |
| Optimized training mixed | Barrera-Domínguez et al. (2020)* | basketball players | -6.69% | -1.67% | 8.45% | 2.11% | -17.82% | -4.45% |
| Optimized training mixed | Barrera-Domínguez et al. (2020)** | basketball players | 6.09% | 1.52% | 7.01% | 1.75% | -0.35% | -0.09% |
| Hypertrophy training | Jonson et al. (2026)* | football players | 0.39% | 0.10% | 12.57% | 3.14% | -12.15% | -3.04% |
| Power training | Jonson et al. (2026)** | football players | 16.60% | 4.15% | 0.24% | 0.06% | 15.42% | 3.85% |

$P_{\max}$ , maximal power output; $wk^{-1}$ = per week (weekly rate of change); $F_0$ , force intercept; $v_0$ , velocity intercept \*First training phase; \*\*Second training phase.

When training is matched to an athlete’s specific F − V_IMB_, both *F*_0_ and *v*_0_improve at a remarkably consistent rate of approximately 2-4% per week (Jiménez-Reyes et al., 2017; Jiménez-Reyes et al., 2019). This finding suggests both variables respond at feasible rates of adaptation across diverse populations. Notably, training one intercept suppresses the other, and this effect is directionally asymmetric: force-targeted interventions substantially reduce *v*_0_ (−10.97% to −33.33%), whereas velocity-targeted interventions attenuate *F*_0_ to a much smaller extent (−3.97% to −13.74%), likely because heavy-load protocols carry a greater velocity cost than ballistic training carries a force cost (Jiménez-Reyes et al., 2017; Jiménez-Reyes et al., 2019). For well-balanced athletes, optimized training produces modest concurrent gains in both variables (1-4% for *F*_0_), although changes in *v*_0_ were less consistent (ranging from −8.20% to +5.59%) (Jiménez-Reyes et al., 2017; Zabaloy et al., 2020). This pattern generally supports a maintenance-oriented rather than a deficit-correction approach. Non-optimized training fails to elicit meaningful *F*_0_ adaptation (−2.64% to +0.63%), yet *v*_0_ improves spontaneously (+1.99% to +10.36%), indicating that velocity capacity might be more readily stimulated by generic athletic training, whereas force capacity requires deliberate, high-load prescription (Barrera-Domínguez et al., 2023; Jiménez-Reyes et al., 2017; Simpson et al., 2021; Zabaloy et al., 2020). Finally, given the limited literature on F-V relationship training, the present discussion is inevitably constrained and should be interpreted with caution.

### Limitations and Future Directions

Several limitations of the present work should be noted. First, the proposed λ coefficient was derived and cross-validated using retrospective datasets drawn from a relatively narrow range of athletic populations; prospective intervention studies with longitudinal monitoring, together with independent validation in female, youth, and elite cohorts, are required to establish its broader generalizability. Future research should also examine how training-induced adaptations and fatigue differentially influence the F-V relationship variables and corresponding elasticity metrics, offering a more comprehensive and integrative framework for individualizing training prescription and monitoring fatigue-related changes over time.

## Conclusion

The present study links distance-averaged velocity to jump height through theoretical derivation and empirical calibration, thereby establishing an elasticity framework for the distance-averaged F-V relationship. This framework quantifies how individual F-V relationship variables govern jump height, with F − V_ER_ indicating which variable dominates the jump height response and F − V_EN_ quantifying the combined jump-height sensitivity to proportional changes in F-V relationship variables. We observed that athletes with higher *F*_0_ or *v*_0_ tend to exhibit correspondingly lower *F*_*e*_ or *v*_*e*_, and that *F*_*e*_ and *v*_*e*_ are inversely coupled. Furthermore, F − V_EN_ and F − V_ER_displayed a U-shaped relationship under the same ℎ_*po*_ and jump height, with experiment showing no straightforward linear association between the two; a balanced profile (F − V_ER_ = 1) did not always correspond to the lowest F − V_EN_. This framework quantifies the jump height response to changes in F-V relationship variables, enabling more precise and individualized training decisions. Together, these findings provide a concise and interpretable tool for informing jump-specific training decisions and assessing athletic potential.

## Acknowledgments

The authors would like to sincerely thank all the participants who volunteered their time and effort to take part in this study.

## Abbreviation

CMJ: countermovement jump
*F̅*_*distance*_: distance-averaged mean force
F-V relationship: force-velocity relationship
F − V_EN_: force-velocity elasticity norm
F − V_ER_: force-velocity elasticity ratio
F − V_IMB_: the ratio between the athlete’s actual F-V slope and the theoretical optimal F-V slope that maximizes jump height at a given maximal power output
*F*_*e*_: force elasticity
*F*_0_: force intercept from F-V relationship
*g*: gravitational acceleration
ℎ: jump height
ℎ_*po*_: push-off distance
*I*(*u*): cumulative net-work integral
*κ*: composite parameter for solving jump height
m: body mass
M: ratio of system mass to body mass
SE: systematic error
SJ: squat jump
*u*: normalized center-of-mass displacement
*v*_*e*_: velocity elasticity
*v̅*_*distance*_: distance-averaged mean velocity
*v*_0_: velocity intercept from F-V relationship
*v*_*to*_: take-off velocity
P_max_: maximal power output
RE: random error
λ: constant used to estimate distance-averaged velocity

