## Appendix1 for "Beyond Imbalance: An Elasticity Framework for the Distance-averaged Force-Velocity Relationship in Vertical Jump"

During the push-off phase, the subject’s acceleration evolves according to the chain rule as follows:

$$\begin{aligned} F\left( x \right)-g=\frac{dv}{dt}=\frac{dv}{dx}\frac{dx}{dt}= v\frac{dv}{dx}=\frac{d{(v}^{2})}{2dx} \#\left( 1 \right) \end{aligned}$$

*F*(*x*) denotes the mass-normalized ground reaction force. Normalizing the ground reaction force to gravity $f\left( x \right) = F(x)/g$, into equation (1) yields:

$$\begin{aligned} 2g\left( f\left( x \right)-1 \right)=\frac{{d(v}^{2})}{dx} \#\left( 2 \right) \end{aligned}$$

Define ${u=x/h}_{po}$∈ [0,1], where $h_{po}$ is vertical displacement of the center of mass from movement onset to take-off. Hence ${dx=h}_{po}du$ and equation becomes

$$\begin{aligned} \frac{{d(v}^{2})}{du}=2gh_{po}\left( f\left( u \right)-1 \right)\#\left( 3 \right) \end{aligned}$$

Integrating with $v(0)=0$,

$$\begin{aligned} v^{2}\left( u \right)=2gh_{po}\int_{0}^{u} \left( f\left( \xi\right)-1 \right) d\xi\#\left( 4 \right) \end{aligned}$$

Define the cumulative net-work integral

$$\begin{aligned} I\left( u \right)=\int_{0}^{u} \left( f\left( \xi\right)-1 \right) d\xi\#\left( 5 \right) \end{aligned}$$

Hence,

$$\begin{aligned} v^{2}\left( u \right)=2g h_{po}I\left( u \right)\#\left( 6 \right) \end{aligned}$$

The $\overline{v}_{\text{distance}}$ is:

$$\begin{aligned} \overline{v}_{\text{distance}}=\int_{0}^{1} v\left( u \right) du=\sqrt{2g h_{po}} \int_{0}^{1} \sqrt{I\left( u \right)} du\#\left( 7 \right) \end{aligned}$$

At take-off, $u=1$:

$$\begin{aligned} v_{to}=v\left( 1 \right)=\sqrt{2g h_{po}\cdot I\left( 1 \right)}\#\left( 8 \right) \end{aligned}$$

The jump height is

$$\begin{aligned} h =\frac{v_{to}^{2}}{2g}{=h}_{po}\cdot I\left( 1 \right)\#\left( 9 \right) \end{aligned}$$

Substituting Equation (9) into Equation (7) and dividing by Equation (8) yields $\lambda$:

$$\begin{aligned} \lambda=\frac{\overline{v}_{\text{distance}}}{v_{to}}=\frac{\int_{0}^{1} \sqrt{I(u)} du}{\sqrt{I(1)}} \#\left( 10 \right) \end{aligned}$$

It is necessary to characterize the force-displacement profile of the vertical jump to solve this formula. Based on Linthorne’s model (2021) of countermovement jump:

$$\begin{aligned} \frac{F(u)}{g}=F\left( 0 \right)\left( 1-u^{A} \right), A>1\#\left( 11 \right) \end{aligned}$$

$F(0)$ denotes the force at the transition from downward to upward movement (bottom position) in the CMJ, while $u$ is the normalized displacement and $A$ controls the shape of force decay during push-off phase. Then, with the dimensionless force defined as $f\left( x \right) = F(x)/g$ yields:

$$\begin{aligned} f\left( u \right)-1=F\left( 0 \right)\left( 1-u^{A} \right)-1\#\left( 12 \right) \end{aligned}$$

Substituting the force model equation (12) into equation (5) and integrating yields the net impulse:

$$\begin{aligned} I_{CMJ}\left( u \right)=F\left( 0 \right)\left( u-\frac{u^{A+1}}{A+1} \right)-u\#\left( 13 \right) \end{aligned}$$

At $u=1$, the ratio of jump height to $h_{po}$ is

$$\begin{aligned} \frac{h}{h_{po}}=I_{CMJ}\left( 1 \right)=F\left( 0 \right)\frac{A}{A+1} -1\#\left( 14 \right) \end{aligned}$$

So,

$$\begin{aligned} F\left( 0 \right)=\left( \frac{h}{h_{po}}+1 \right)\frac{A+1}{A}\#\left( 15 \right) \end{aligned}$$

Substituting into equation (15):

$$\begin{aligned} \lambda=\frac{\int_{0}^{1} \sqrt{F(0)\left( u-\frac{u^{A+1}}{A+1} \right)-u} du}{\sqrt{F(0)\frac{A}{A+1}-1}}\#\left( 16 \right) \end{aligned}$$

For the typical ranges of $F(0)$ (range from 2 to 3.5) and $A$ (range from 3 to 5), the computed $\lambda$ mean value is 0.77 (SD = 0.02, 95% CI [0.75, 0.83]). This result is consistent with the $\lambda$ values obtained from re-analyzing our previous squat jump data within the same framework, which also yields $\lambda$ ≈0.77.
