## Appendix2 for "Beyond Imbalance: An Elasticity Framework for the Distance-averaged Force-Velocity Relationship in Vertical Jump"

The ${F-V}_{\mathrm{IMB}}$ is defined as the slope ratio:

$$\begin{aligned} {F-V}_{\mathrm{IMB}}=\frac{S}{S_{\mathrm{opt}}}\#\left( 1 \right) \end{aligned}$$

where $S=F_{0}/v_{0}$ is the slope of the linear F-V relationship and $S_{\mathrm{opt}}$ is the slope that maximises jump height for a given $P_{\max}$. The ${F-V}_{\mathrm{ER}}$ is defined as:

$$\begin{aligned} {F-V}_{\mathrm{ER}}=\frac{F_{e}}{v_{e}}\#\left( 2 \right) \end{aligned}$$

The elasticities of jump height with respect to the $F_{0}$ and $v_{0}$ admit the closed forms:

$$\begin{aligned} F_{e}=\frac{2F_{0} (v_{0}-\lambda\sqrt{2gh})}{\frac{2gh v_{0}}{h_{po}}+F_{0} \lambda\sqrt{2gh}}\#\left( 3 \right) \end{aligned}$$

$$\begin{aligned} v_{e}=\frac{2F_{0} \lambda\sqrt{2gh}}{\frac{2gh v_{0}}{h_{po}}+F_{0} \lambda\sqrt{2gh}}\#\left( 4 \right) \end{aligned}$$

Dividing equation (3) by equation (4), the common denominator cancels:

$$\begin{aligned} {F-V}_{\mathrm{ER}}=\frac{F_{e}}{v_{e}}=\frac{v_{0}}{\lambda\sqrt{2gh}}-1\#\left( 5 \right) \end{aligned}$$

The mean force over the push-off is $\bar{F}_{distance}=F_{0}-S \lambda\sqrt{2gh}$, and the jump height follows from the work-energy balance:

$$\begin{aligned} h=\frac{h_{po}}{g}\left( F_{0}-S \lambda\sqrt{2gh}-g \right)\#\left( 6 \right) \end{aligned}$$

Fix the maximal power output $P_{\max}=F_{0}v_{0}/4$, i.e. $F_{0}=2\sqrt{P_{\max}S}$. The optimal slope maximises $h$; differentiating h with respect to $S$ and setting $\partial h/\partial S=0$ gives:

$$\begin{aligned} S_{\mathrm{opt}}=\frac{P_{\max}}{2\lambda^{2}gh} (6)\#\left( 7 \right) \end{aligned}$$

Substituting equation (7) into the equation (1):

$$\begin{aligned} {F-V}_{\mathrm{IMB}}=\frac{S}{S_{\mathrm{opt}}}=\frac{F_{0}/v_{0}}{P_{\max}/(2\lambda^{2}gh)}=\frac{8\lambda^{2}gh}{v_{0}^{2}}\#\left( 8 \right) \end{aligned}$$

From equation (5), $v_{0}=\lambda\sqrt{2gh} ({F-V}_{\mathrm{ER}}+1)$; hence $v_{0}^{2}=2\lambda^{2}gh ({F-V}_{\mathrm{ER}}+1)^{2}$. Substituting into equation (8):

$$\begin{aligned} {F-V}_{\mathrm{IMB}}=\frac{4}{({F-V}_{\mathrm{ER}}+1)^{2}}\#\left( 9 \right) \end{aligned}$$

and inverting:

$$\begin{aligned} {F-V}_{\mathrm{ER}}=\frac{2}{\sqrt{{F-V}_{\mathrm{IMB}}}}-1\#\left( 10 \right) \end{aligned}$$

A balanced F-V relationship (${F-V}_{\mathrm{IMB}}=1$, where the actual slope equals the optimal slope) corresponds to ${F-V}_{\mathrm{ER}}=1$, indicating equal $F_{e}$ and $v_{e}$; the two conditions are equivalent.
