## Appendix3 for "Beyond Imbalance: An Elasticity Framework for the Distance-averaged Force-Velocity Relationship in Vertical Jump"

The derivation begins with the distance-averaged force–velocity framework and the elasticity definitions:

$$\begin{aligned} \bar{v}_{distance}=\lambda\sqrt{2gh}\#\left( 1 \right) \end{aligned}$$

$$\begin{aligned} \bar{F}_{distance}=\frac{g(h_{po}+h)}{h_{po}} \#\left( 2 \right) \end{aligned}$$

$$\begin{aligned} \bar{F}_{distance}=F_{0}-\frac{F_{0}}{v_{0}}\bar{v}_{distance} \#\left( 3 \right) \end{aligned}$$

$$\begin{aligned} F_{e}=\frac{\%\Delta h}{\%\Delta F_{0}}=\frac{\partial(\ln h)}{\partial(\ln F_{0})} \#\left( 4 \right) \end{aligned}$$

$$\begin{aligned} v_{e}=\frac{\%\Delta h}{\%\Delta v_{0}}=\frac{\partial(\ln h)}{\partial(\ln v_{0})} \#\left( 5 \right) \end{aligned}$$

Substituting (1) and (2) into (3) eliminates $\bar{v}_{\mathrm{distance}}$ and $\bar{F}_{\mathrm{distance}}$:

$$\begin{aligned} F_{0}-\frac{F_{0}}{v_{0}} \lambda\sqrt{2gh} = g+\frac{g h}{h_{po}} \#\left( 6 \right) \end{aligned}$$

Rearranging defines an implicit function $\Phi(h,F_{0},v_{0})=0$:

$$\begin{aligned} \Phi: F_{0}-\frac{F_{0}\lambda}{v_{0}}\sqrt{2gh}-g-\frac{g h}{h_{po}}=0 \#\left( 7 \right) \end{aligned}$$

Here $\Phi$ is a function of $h$,$F_{0}$, and $v_{0}$, where $F_{0}$ and $v_{0}$ are independent variables, and $h$ is a function of$F_{0}$ and $v_{0}$:$h=h(F_{0},v_{0})$. Partial derivatives via implicit differentiation

For $\Phi(h,F_{0},v_{0})=0$,

$$\begin{aligned} \frac{\partial h}{\partial F_{0}}=-\frac{{\partial\Phi}/{\partial F_{0}}}{{\partial\Phi}/{\partial h}} \#\left( 8 \right) \end{aligned}$$

$$\begin{aligned} \frac{\partial h}{\partial v_{0}}=-\frac{{\partial\Phi}/{\partial v_{0}}}{{\partial\Phi}/{\partial h}} \#\left( 9 \right) \end{aligned}$$

Computing each partial derivative of $\Phi$:

$$\begin{aligned} \frac{\partial\Phi}{\partial h}=-\frac{F_{0}\lambda\sqrt{2g}}{2v_{0}\sqrt{h}}-\frac{g}{h_{po}} \#\left( 10 \right) \end{aligned}$$

$$\begin{aligned} \frac{\partial\Phi}{\partial F_{0}}=1-\frac{\lambda\sqrt{2gh}}{v_{0}} \#\left( 11 \right) \end{aligned}$$

$$\begin{aligned} \frac{\partial\Phi}{\partial v_{0}}=\frac{F_{0}\lambda\sqrt{2gh}}{v_{0}^{2}} \#\left( 12 \right) \end{aligned}$$

Hence

$$\begin{aligned} \frac{\partial h}{\partial F_{0}}=\frac{1-\frac{\lambda\sqrt{2gh}}{v_{0}}}{\frac{F_{0}\lambda\sqrt{2g}}{2v_{0}\sqrt{h}}+\frac{g}{h_{po}}} \#\left( 13 \right) \end{aligned}$$

$$\begin{aligned} \frac{\partial h}{\partial v_{0}}=\frac{\frac{F_{0}\lambda\sqrt{2gh}}{v_{0}^{2}}}{\frac{F_{0}\lambda\sqrt{2g}}{2v_{0}\sqrt{h}}+\frac{g}{h_{po}}} \#\left( 14 \right) \end{aligned}$$

Expressions for $F_{e}$ and $v_{e}$

Inserting the partial derivatives into (4) and (5):

$$\begin{aligned} F_{e}=\frac{F_{0}}{h} \frac{1-\frac{\lambda\sqrt{2gh}}{v_{0}}}{\frac{F_{0}\lambda\sqrt{2g}}{2v_{0}\sqrt{h}}+\frac{g}{h_{po}}} \#\left( 15 \right) \end{aligned}$$

$$\begin{aligned} v_{e} =\frac{F_{0}\lambda\sqrt{2gh}}{v_{0} h} \frac{1}{\frac{F_{0}\lambda\sqrt{2g}}{2v_{0}\sqrt{h}}+\frac{g}{h_{po}}} \#\left( 16 \right) \end{aligned}$$

Both share the same positive denominator; therefore $F_{e}=v_{e}$ reduces to equality of their numerators.

Balance Condition $F_{e}=v_{e}$

Cancelling the common positive factor $F_{0}/h$,

$$\begin{aligned} 1-\frac{\lambda\sqrt{2gh}}{v_{0}}=\frac{\lambda\sqrt{2gh}}{v_{0}} \#\left( 17 \right) \end{aligned}$$

Simplifying,

$$\begin{aligned} v_{0}=2\lambda\sqrt{2gh} \#\left( 18 \right) \end{aligned}$$

Squaring and solving for $h$:

$$\begin{aligned} h=\frac{v_{0}^{2}}{8\lambda^{2}g} \#\left( 19 \right) \end{aligned}$$

At $F_{e}=v_{e}$, jump height is determined solely by $v_{0}$

Returning to equation (7) and substituting equation (19):

$$\begin{aligned} \frac{F_{0}}{2}-g=\frac{g}{h_{po}}\cdot\frac{v_{0}^{2}}{8\lambda^{2}g} \#\left( 20 \right) \end{aligned}$$

Hence,

$$\begin{aligned} F_{0}=2g+\frac{v_{0}^{2}}{4\lambda^{2}h_{po}}\#\left( 21 \right) \end{aligned}$$

Equation (21) defines the locus of $(F_{0},v_{0})$ pairs at which $F_{e}=v_{e}$ for a given $h_{po}$. Taking the differential of both sides and simplifying yield:

$$\begin{aligned} \frac{dF_{0}}{F_{0}-2g}=2 \frac{dv_{0}}{v_{0}}\#\left( 22 \right) \end{aligned}$$

Notably, this relationship is independent of $h_{po}$, $\lambda$ and $g$.
